# Beware of White Matter Hyperintensities Causing Systematic Errors in Grey Matter Segmentations!

**DOI:** 10.1101/2020.07.07.191809

**Authors:** Mahsa Dadar, Olivier Potvin, Richard Camicioli, Simon Duchesne, for the Alzheimer’s Disease Neuroimaging Initiative

**Affiliations:** CERVO Brain Research Center, Centre intégré universitaire santé et services sociaux de la Capitale Nationale, Québec, QC; Department of Medicine, Division of Neurology, University of Alberta, Edmonton, AB; Department of Radiology and Nuclear Medicine, Faculty of Medicine, Université Laval, Québec, QC

**Keywords:** White matter hyperintensities, grey matter segmentation, FreeSurfer, Alzheimer’s disease

## Abstract

**Introduction:** Volumetric estimates of subcortical and cortical structures, extracted from T1-weighted MRIs, are widely used in many clinical and research applications. Here, we investigate the impact of the presence of white matter hyperintensities (WMHs) on *FreeSurfer* grey matter (GM) structure volumes and its possible bias on functional relationships.

**Methods:** T1-weighted images from 1077 participants (4321 timepoints) from the Alzheimer’s Disease Neuroimaging Initiative were processed with *FreeSurfer* version 6.0.0. WMHs were segmented using a previously validated algorithm on either T2-weighted or Fluid-attenuated inversion recovery (FLAIR) images. Mixed effects models were used to assess the relationships between overlapping WMHs and GM structure volumes and overal WMH burden, as well as to investigate whether such overlaps impact associations with age, diagnosis, and cognitive performance.

**Results:** Participants with higher WMH volumes had higher overalps with GM volumes of bilateral caudate, cerebral cortex, putamen, thalamus, pallidum, and accumbens areas (P < 0.0001). When not corrected for WMHs, caudate volumes increased with age (P < 0.0001) and were not different between cognitively healthy individuals and age-matched probable Alzheimer’s disease patients. After correcting for WMHs, caudate volumes decreased with age (P < 0.0001), and Alzheimer’s disease patients had lower caudate volumes than cognitively healthy individuals (P < 0.01). Uncorrected caudate volume was not associated with ADAS13 scores, whereas corrected lower caudate volumes were significantly associated with poorer cognitive performance (P < 0.0001).

**Conclusions:** Presence of WMHs leads to systematic inaccuracies in GM segmentations, particularly for the caudate, which can also change clinical associations. While specifically measured for the *Freesurfer* toolkit, this problem likely affects other algorithms.

## INTRODUCTION

White matter hyperintenesities (WMHs) are defined as areas of increased signal on T2-weighted (T2w) and Fluid-attenuated inversion recovery (FLAIR) magnetic resonance images (MRIs) (Raman et al., 2016). WMHs are associated with a variety of underlying pathologies, such as amyloid angiopathy, arteriosclerosis, axonal loss, blood-brain barrier leakage, degeneration, demyelination, gliosis, hypoperfusion, hypoxia, and inflammation (Abraham et al., 2016). WMHs are commonly present in the otherwise asymptomatic aging population but are at higher prevalence in many diseases such as Alzheimer’s disease (AD), diabetes, frontotemporal dementia, HIV, lewy body dementia, mild cognitive impairment (MCI), obesity, Parkinson’s disease, and vascular dementia (Appelman et al., 2009; Debette and Markus, 2010; Caroppo et al., 2014; Dadar et al., 2018b; Gouw et al., 2008; Kandiah et al., 2013; Sudre et al., 2017; Barber et al., 1999).

On T1-weighted (T1w) MRI sequences, WMHs appear hypointense with respect to the normal-appearing white matter, with intensities that can be very similar to cortical and subcortical grey matter (GM) (Dadar et al., 2017a). The T1w intensity of WMHs is also associated with severity of damage to the tissue, with areas of higher damage appearing more hypointense (Dadar et al., 2019).

As T1w images are the most commonly used structural MRI sequences in clinical and neuroscience applications, especially for purposes of segmentation and estimation of volumes for all or specific structures of interest (Mateos-Pérez et al., 2018), the similarity in T1w intensity profiles of WMHs and GM gives rise to an important methodological question: can T1w MRI-based GM structure segmentation differentiate between WMHs and GM? If not, how much of WMHs will be tagged as GM in their segmentation estimates, and if so, is this error systematic enough to bias results?

To answer this question, we propose our study of WMHs segmentation bias in subcortical and cortical GM structures. More specifically, we investigated 1) whether there was any systematic overlap between WMHs and GM segmentations, and therefore volumetric biases; and 2) whether this overlap affected clinical findings (i.e. associations with cognitive scores). To these ends we used two segmentation tools, the first being *FreeSurfer,* one of the most commonly used publicly available brain segmentation tools (Fischl, 2012), and a previously validated tool for WMHs segmentation on multi-contrast MRIs (Dadar et al., 2017b), both applied on longitudinal data from the Alzheimer’s Disease Neuroimaging Initiative (ADNI) database.

## METHODS

### Participants

We used longitudinal data from 1077 participants (4321 individual images from various timepoints) from the ADNI-1, ADNI-2, and ADNI-GO database (adni.loni.usc.edu) that had T1w and either T2w/PDw or FLAIR MRIs available. The ADNI was launched in 2003 as a public-private partnership, led by Principal Investigator Michael W. Weiner, MD. The primary goal of ADNI has been to test whether serial MRI, positron emission tomography, other biological markers, and clinical and neuropsychological assessment can be combined to measure the progression of MCI and early AD. The study was approved by the institutional review board of all participating sites and written informed consent was obtained from all participants before inclusion in the study.

### MRI acquisition and preprocessing

Table 1 summarizes MR imaging parameters for the data used in this study.

**Table 1.** Scanner information and MRI acquisition parameters for ADNI1, and ADNI2/GO datasets.

|  | Sequence | T1w | T2w/PDw | FLAIR |
| --- | --- | --- | --- | --- |
| ADNI1 | Slice thickness | 1.2 mm | 3 mm |  |
|  | No. of slices | 160 | 56 |  |
|  | Field of view | 192×192 cm<sup>2</sup> | 256×256 cm<sup>2</sup> |  |
|  | Scan Matrix | 192×192 cm<sup>2</sup> | 256×256 cm<sup>2</sup> |  |
|  | Repetition time (TR) | 3000 ms | 3000/3000 ms |  |
|  | Echo time (TE) | 3.55 ms | 95.2/10.5 ms |  |
| ADNI2/GO | Slice thickness | 1.2 mm |  | 5 mm |
|  | No. of slices | 196 |  | 42 |
|  | Field of view | 256×256 cm<sup>2</sup> |  | 256×256 cm<sup>2</sup> |
|  | Scan Matrix | 256×256 cm<sup>2</sup> |  | 256×256 cm<sup>2</sup> |
|  | Repetition time (TR) | 7.2 ms |  | 11000 ms |
|  | Echo time (TE) | 3.0 ms |  | 150 ms |

### GM Segmentations

All T1w images were identically processed using *FreeSurfer* version 6.0.0 *(recon-all-all). FreeSurfer* is an open source software (https://surfer.nmr.mgh.harvard.edu/) that provides a full processing stream for structural T1w data (Fischl, 2012). The final segmentation output (aseg.mgz) was then used to obtain structure masks and volumes based on the look up table available at https://surfer.nmr.mgh.harvard.edu/fswiki/FsTutorial/AnatomicalROI/FreeSurferColorLUT.

### WMH Segmentations

T1w, T2w/PDw, and FLAIR scans were pre-processed as follows: (a) image denoising (Manjón et al., 2010); (b) intensity inhomogeneity correction; and (c) intensity normalization to a 0-100 range. For each subject, the T2w, PDw, or FLAIR scans were then co-registered to the structural T1w scan of the same timepoint using a 6-parameter rigid registration and a mutual information objective function (Dadar et al., 2018a). We used a previously validated WMH segmentation method that employs a set of location and intensity features in combination with a random forests classifier to detect WMHs using either T1w+FLAIR or T1w+T2w/PDw images. The training library consisted in manual, expert segmentations of WMHs from 100 subjects from ADNI (not included in the current sample) (Dadar et al., 2017c, 2017b, 2018b). WMHs were automatically segmented at all timepoints and then co-registered to the T1w images using the obtained rigid registrations, in order to assess overlaps between WMHs and GM segmentations.

### Cognitive Evaluations

All subjects received a comprehensive battery of clinical assessments and cognitive testing based on a standardized protocol (adni.loni.usc.edu) (Petersen et al., 2010). At each visit, participants underwent a series of assessments including the Alzheimer’s Disease Assessment Scale-13 (ADAS13) (Mohs and Cohen, 1987), which was used to assess cognitive performance.

### Statistical Analyses

Overlaps between GM and WMH segmentations were calculated (number of overlapping voxels in mm^3^) for each subcortical and cortical GM region. The following mixed effects models were used to assess whether the WMH-GM overlaps were associated with overall WMH burden, controlling for age and sex.

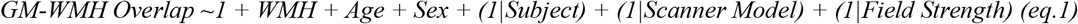

Mixed effects models were also used to assess the relationships between GM volumes and age and diagnostic cohort, and GM volumes and cognition, once using the GM volume estimates obtained directly from the FreeSurfer segmentation, and once after removing the regions overlapping with the WMH segmentations.

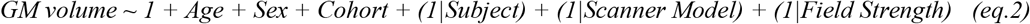

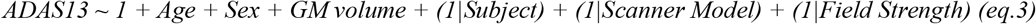

All volumes were normalized by the individual’s intracranial volume. Total WMH loads and WMH-GM overlaps were log-transformed to obtain normal distributions. All mixed effects models included Subject as well as Scanner Model and Field Strength as categorical random variables, to account for any potential variabilities caused by contrast differences in the images from different scanners. The results were corrected for multiple comparisons using the false discovery rate (FDR) controlling method with a significance threshold of P = 0.05 (Benjamini and Hochberg, 1995).

### Data and Code Availability Statement

Data used in this article is available at http://adni.loni.usc.edu/. FreeSurfer and the WMH segmentation pipeline used are also publicly available at https://surfer.nmr.mgh.harvard.edu/ and http://nist.mni.mcgill.ca/?p=221, respectively.

## RESULTS

### Study participants

Preprocessed and registered images were visually assessed for quality control (presence of imaging artifacts, failure in registrations). WMH segmentations were also visually assessed for missing hyperintensities or over-segmentation. Either failures resulted in the participant being removed from the analyses. All MRI processing, segmentation and quality control steps were blinded to clinical outcomes. All cases passed co-registration QC. Figure 1 summarizes the QC information for the subjects that were excluded. The final sample included 1077 participants (4321 timpoints) with WMH and FreeSurfer segmentations.

**Figure 1.**
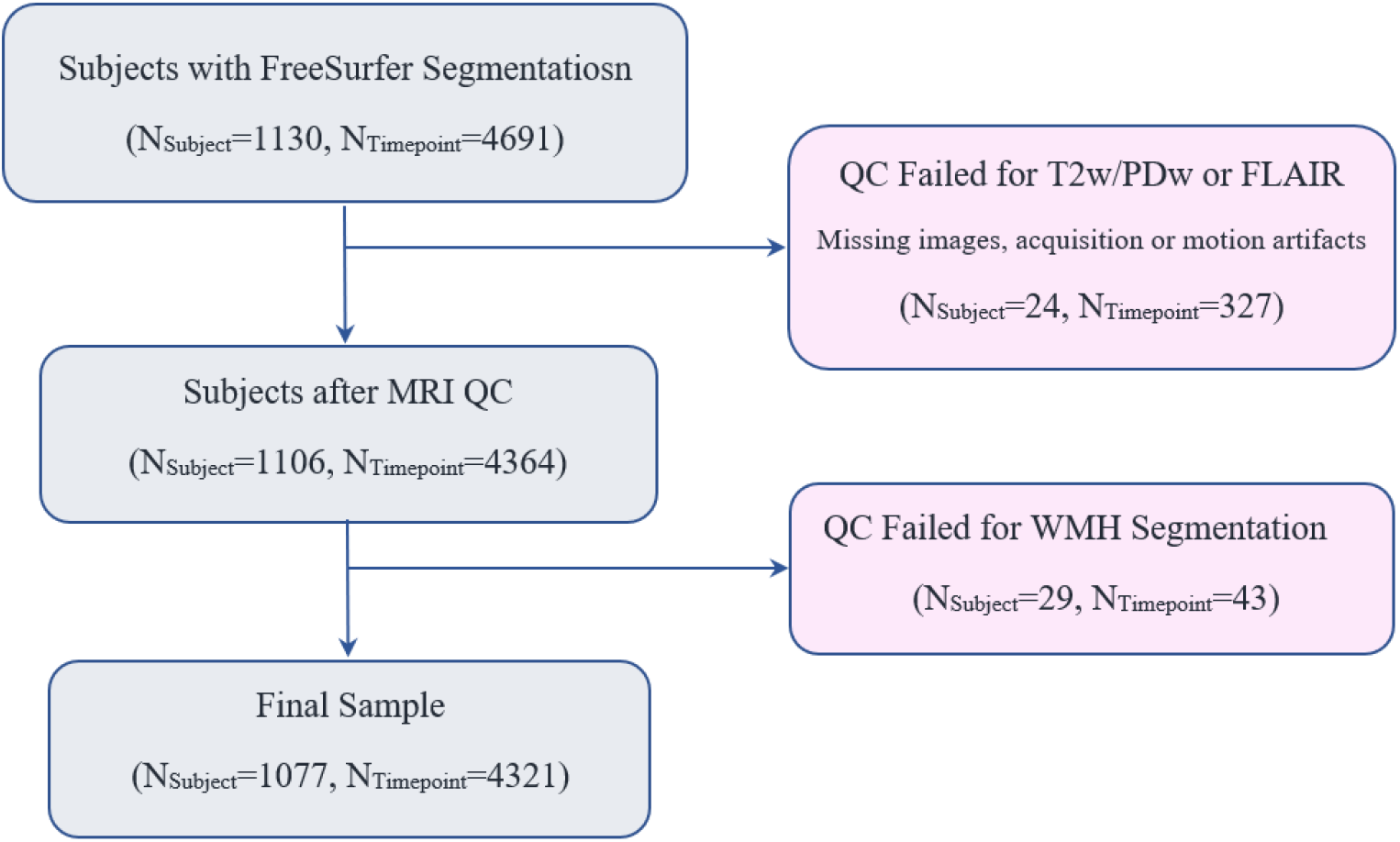
Flowchart of subjects in the study.

### Segmentations

Figure 2 shows WMH and caudate segmentations for three subjects with no overlap, some overlap, and high overlap as an example.

**Figure 2.**
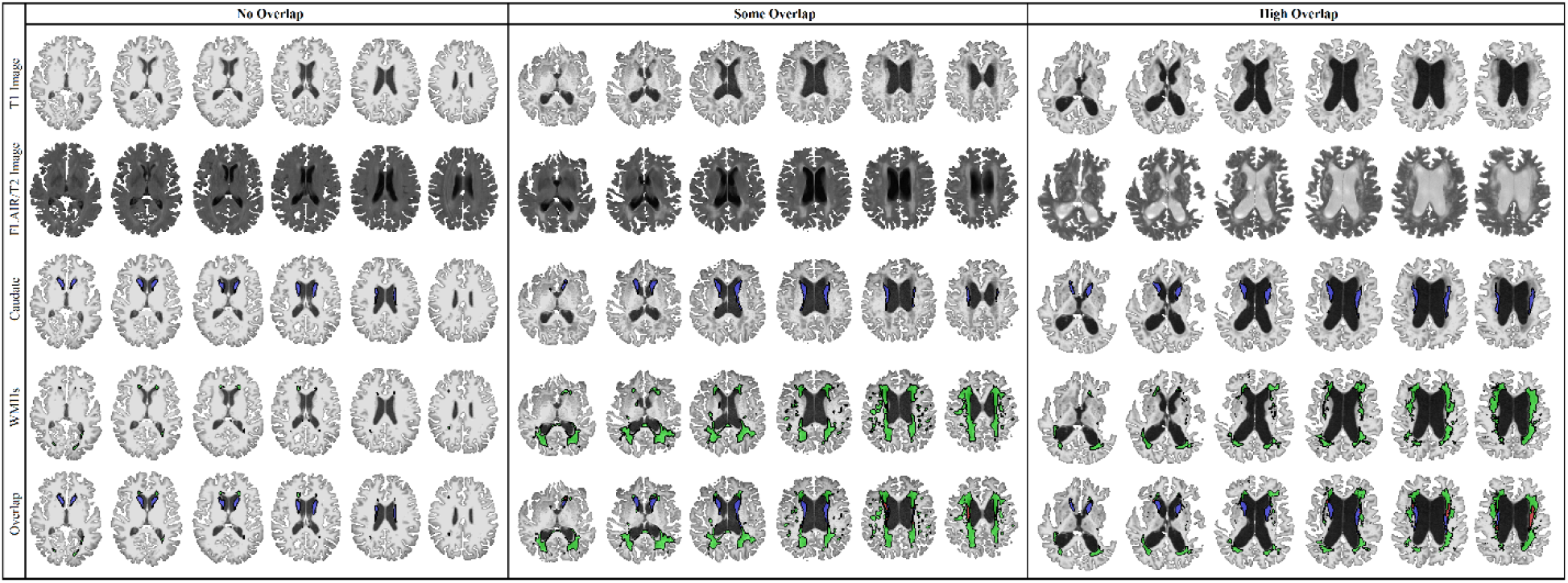
Overlap between WMH and caudate segmentations. WMH= White Matter Hyperintensity. Blue= Caudate. Green= WMHs. Red= The overlapping voxels between caudate and WMH segmentations.

### Segmentation overlap

Table 2 shows the average amount of overlap between WMHs and FreeSurfer segmentations for each GM structure, as well as their association with the overall WMH burden, controlling for age and sex (eq.1). Caudate segmentations had by far the highest percentage of overlapping WMHs (6% of the mean caudate volume). The overlapping volumes were significantly related to the overall WMH burden for bilateral caudate, cerebral cortex, putamen, thalamus, pallidum, accumbens area, and the right hippocampus (P<0.0002). Figure 2 shows the associations for the top six regions.

**Table 2.** Overlaps between WMHs and FreeSurfer GM segmentations, and their associations with overal WMH burden. The regions are sorted based on effect size. Significant results after FDR correction are indicated in bold font.

| Structure Name | FreeSurfer Label | Volume (mm<sup>3</sup>) | Overlap with WMH (mm<sup>3</sup>) | Percentage of Overlap (%) | Association with WMH Tstat | Pvalue |
| --- | --- | --- | --- | --- | --- | --- |
| Caudate - Right | 50 | 3532.9 ± 596.2 | 217.2 ± 271.0 | 6.153 | **45.80** | **<0.0001** |
| Caudate - Left | 11 | 3395.9 ± 556.4 | 222.9 ± 283.9 | 6.565 | **45.21** | **<0.0001** |
| Cerebral Cortex - Left | 3 | 202864 ± 25017 | 73.8 ± 240.5 | 0.036 | **31.41** | **<0.0001** |
| Cerebral Cortex - Right | 42 | 203763 ± 25182 | 79.8 ± 218.8 | 0.039 | **29.54** | **<0.0001** |
| Putamen - Right | 51 | 4235.6 ± 644.1 | 21.9 ± 74.4 | 0.518 | **22.77** | **<0.0001** |
| Putamen - Left | 12 | 4194.8 ± 648.3 | 17.4 ± 56.1 | 0.416 | **20.88** | **<0.0001** |
| Thalamus - Left | 10 | 6598.1 ± 729.3 | 1.01 ± 5.72 | 0.015 | **11.94** | **<0.0001** |
| Thalamus - Right | 49 | 6499.9 ± 711.3 | 0.55 ± 4.07 | 0.008 | **8.53** | **<0.0001** |
| Pallidum - Right | 52 | 1829.0 ± 256.6 | 0.85 ± 3.50 | 0.047 | **8.52** | **<0.0001** |
| Accumbens Area - Left | 26 | 405.1 ± 86.6 | 0.23 ± 1.96 | 0.057 | **7.67** | **<0.0001** |
| Pallidum - Left | 13 | 1863.1 ± 255.1 | 0.45 ± 2.50 | 0.024 | **5.70** | **<0.0001** |
| Accumbens Area - Right | 58 | 452.7 ± 90.7 | 0.17 ± 0.88 | 0.038 | **4.67** | **<0.0001** |
| Hippocampus - Right | 53 | 3572.6 ± 600.3 | 0.22 ± 2.24 | 0.006 | **3.72** | **0.0002** |
| Ventral Diencephalon - Right | 60 | 3754.9 ± 459.5 | 0.10 ± 1.78 | 0.003 | 1.86 | 0.10 |
| Amygdala - Right | 54 | 1469.7 ± 319.3 | 0.04 ± 0.88 | 0.003 | 1.58 | 0.11 |
| Amygdala - Left | 18 | 1275.3 ± 299.1 | 0.01 ± 0.23 | 0.0001 | 0.70 | 0.48 |
| Ventral Diencephalon - Left | 28 | 3782.2 ± 473.9 | 0.06 ± 2.03 | 0.002 | 0.69 | 0.48 |
| Hippocampus - Left | 17 | 3476.3 ± 564.9 | 0.17 ± 4.77 | 0.005 | 0.36 | 0.71 |

To better demonstrate how much the misclassification of WMHs as caudate might increase caudate volume estimates, we have also plotted volume overlaps for caudate without log-transformation (i.e. volumes in mm^3^) in Figure 4. In extreme cases, for individuals with very high WMH burden (log transformed value of 11, equivalent to 50,000 mm^3^), caudate volume can be over-estimated by more than 1000 mm^3^, equivalent to 30% of the average caudate volume.

**Figure 3.**
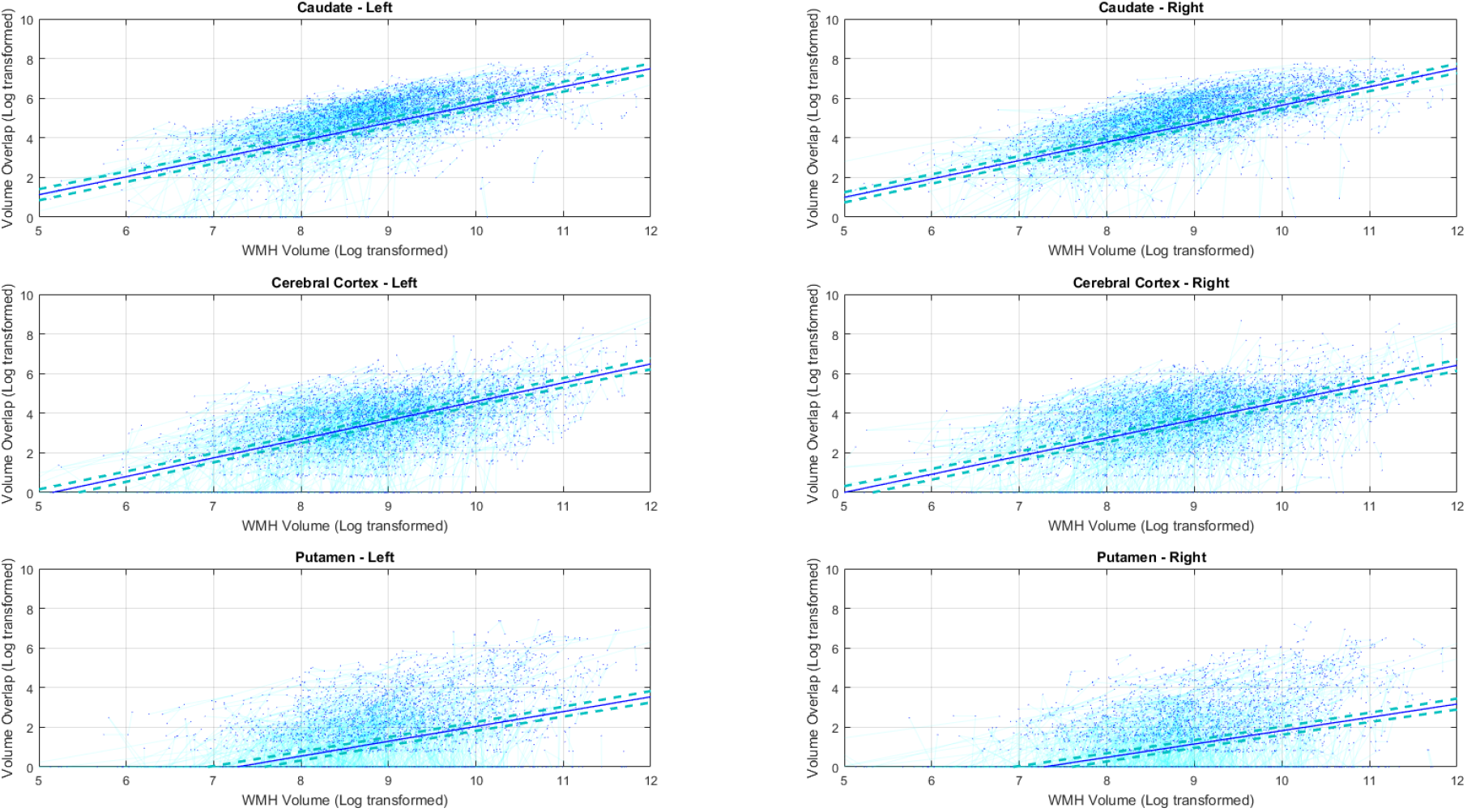
The association between overlapping GM and WMH volumes and overall WMH burden (Table 2).

**Figure 4.**
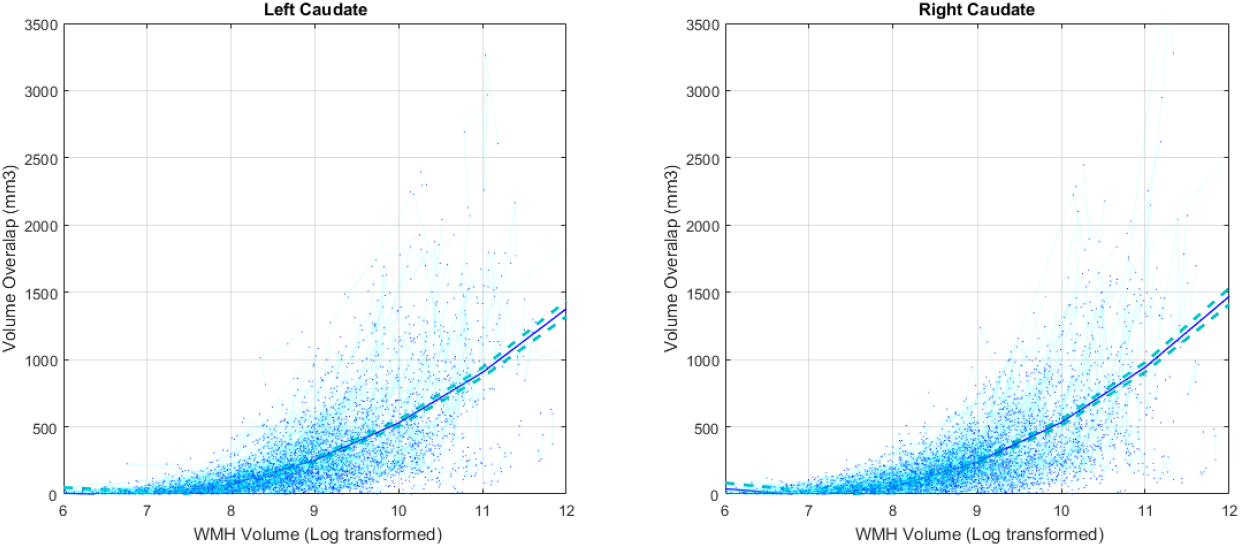
The association between overlapping Caudate and WMH volumes (in mm^3^) and overall WMH burden.

### Associations

Table 3 shows the associations between GM volumes and age (eq. 2), before and after removing the voxels overlapping with WMHs. Uncorrected caudate volumes increased with age, and MCI and AD groups had slightly higher volumes than the normal aging (NA) cohort (although not significant), whereas the corrected caudate volumes decreased with age, and MCI and AD groups had slightly lower volumes than the NA group (the AD vs NA difference was significant). The uncorrected and corrected results were similar in terms of effect size and direction of associations for other regions, with the corrected volumes having slightly larger effects sizes for putamen.

**Table 3.** Associations between uncorrected and corrected GM volumes, age, and diagnostic cohort. Significant results after FDR correction are indicated in bold font.

| Structure Name |  | Age |  | MCI vs NA |  | AD vs NA |  |
| --- | --- | --- | --- | --- | --- | --- | --- |
|  |  | Tstat | Pvalue | Tstat | Pvalue | Tstat | Pvalue |
| Caudate - Right | Uncorrected | **4.24** | **<0.0001** | 0.61 | 0.54 | 0.58 | 0.56 |
|  | Corrected | **-3.77** | **<0.0001** | -0.94 | 0.34 | **-2.46** | **0.01** |
| Caudate - Left | Uncorrected | **5.45** | **<0.0001** | 0.97 | 0.33 | 0.37 | 0.71 |
|  | Corrected | **-6.30** | **<0.0001** | -1.19 | 0.23 | **-2.60** | **0.009** |
| Cerebral Cortex - Left | Uncorrected | **-32.59** | **<0.0001** | **-6.39** | **<0.0001** | **-12.14** | **<0.0001** |
|  | Corrected | **-32.57** | **<0.0001** | **-6.41** | **<0.0001** | **-12.18** | **<0.0001** |
| Cerebral Cortex - Right | Uncorrected | **-32.41** | **<0.0001** | **-6.21** | **<0.0001** | **-11.33** | **<0.0001** |
|  | Corrected | **-32.42** | **<0.0001** | **-6.22** | **<0.0001** | **-11.36** | **<0.0001** |
| Putamen - Right | Uncorrected | **-18.74** | **<0.0001** | **-4.57** | **<0.0001** | **-5.88** | **<0.0001** |
|  | Corrected | **-19.46** | **<0.0001** | **-4.79** | **<0.0001** | **-6.37** | **<0.0001** |
| Putamen - Left | Uncorrected | **-15.56** | **<0.0001** | **-4.61** | **<0.0001** | **-7.19** | **<0.0001** |
|  | Corrected | **-16.30** | **<0.0001** | **-4.84** | **<0.0001** | **-7.59** | **<0.0001** |
| Thalamus - Left | Uncorrected | **-18.84** | **<0.0001** | **-4.93** | **<0.0001** | **-7.52** | **<0.0001** |
|  | Corrected | **-18.84** | **<0.0001** | **-4.94** | **<0.0001** | **-7.52** | **<0.0001** |
| Thalamus - Right | Uncorrected | **-18.68** | **<0.0001** | **-4.10** | **<0.0001** | **-6.92** | **<0.0001** |
|  | Corrected | **-18.69** | **<0.0001** | **-4.09** | **<0.0001** | **-6.92** | **<0.0001** |
| Pallidum - Right | Uncorrected | **-12.75** | **<0.0001** | **-2.50** | **0.01** | **-2.43** | **0.01** |
|  | Corrected | **-12.81** | **<0.0001** | **-2.51** | **0.01** | **-2.45** | **0.01** |
| Accumbens Area - Left | Uncorrected | **-19.77** | **<0.0001** | **-4.08** | **<0.0001** | **-7.10** | **<0.0001** |
|  | Corrected | **-19.83** | **<0.0001** | **-4.08** | **<0.0001** | **-7.11** | **<0.0001** |
| Pallidum - Left | Uncorrected | **-13.76** | **<0.0001** | **-2.25** | **0.02** | **-2.71** | **0.006** |
|  | Corrected | **-13.78** | **<0.0001** | **-2.25** | **0.02** | **-2.72** | **0.006** |
| Accumbens Area - Right | Uncorrected | **-19.77** | **<0.0001** | **-3.32** | **0.0002** | **-6.41** | **<0.0001** |
|  | Corrected | **-19.83** | **<0.0001** | **-3.34** | **0.0002** | **-6.42** | **<0.0001** |
| Hippocampus - Right | Uncorrected | **-33.26** | **<0.0001** | **-11.32** | **<0.0001** | **-16.27** | **<0.0001** |
|  | Corrected | **-33.27** | **<0.0001** | **-11.32** | **<0.0001** | **-16.27** | **<0.0001** |
| Ventral Diencephalon - Right | Uncorrected | **-25.30** | **<0.0001** | **-5.39** | **<0.0001** | **-6.76** | **<0.0001** |
|  | Corrected | **-25.31** | **<0.0001** | **-5.39** | **<0.0001** | **-6.76** | **<0.0001** |
| Amygdala - Right | Uncorrected | **-23.92** | **<0.0001** | **-10.12** | **<0.0001** | **-15.30** | **<0.0001** |
|  | Corrected | **-23.93** | **<0.0001** | **-10.12** | **<0.0001** | **-15.30** | **<0.0001** |
| Amygdala - Left | Uncorrected | **-27.29** | **<0.0001** | **-9.85** | **<0.0001** | **-16.27** | **<0.0001** |
|  | Corrected | **-27.29** | **<0.0001** | **-9.85** | **<0.0001** | **-16.28** | **<0.0001** |
| Ventral Diencephalon - Left | Uncorrected | **-25.23** | **<0.0001** | **-5.24** | **<0.0001** | **-7.54** | **<0.0001** |
|  | Corrected | **-25.23** | **<0.0001** | **-5.23** | **<0.0001** | **-7.54** | **<0.0001** |
| Hippocampus - Left | Uncorrected | **-32.25** | **<0.0001** | **-11.1** | **<0.0001** | **-16.92** | **<0.0001** |
|  | Corrected | **-32.25** | **<0.0001** | **-11.1** | **<0.0001** | **-16.92** | **<0.0001** |

Table 4 shows the associations between ADAS13 and GM volumes, before and after removing the voxels overlapping with WMHs. Uncorrected caudate volumes were not significantly associated with ADAS13 scores, whereas lower corrected caudate volumes were significantly associated with higher ADAS13 scores (i.e. poorer cognitive performance). Figure 5 shows the associations between uncorrected and corrected caudate volumes and ADAS13 scores. The uncorrected and corrected results were similar in terms of effect size and direction of associations for other regions, with the corrected volumes having slightly larger effects sizes for putamen.

**Table 4.**
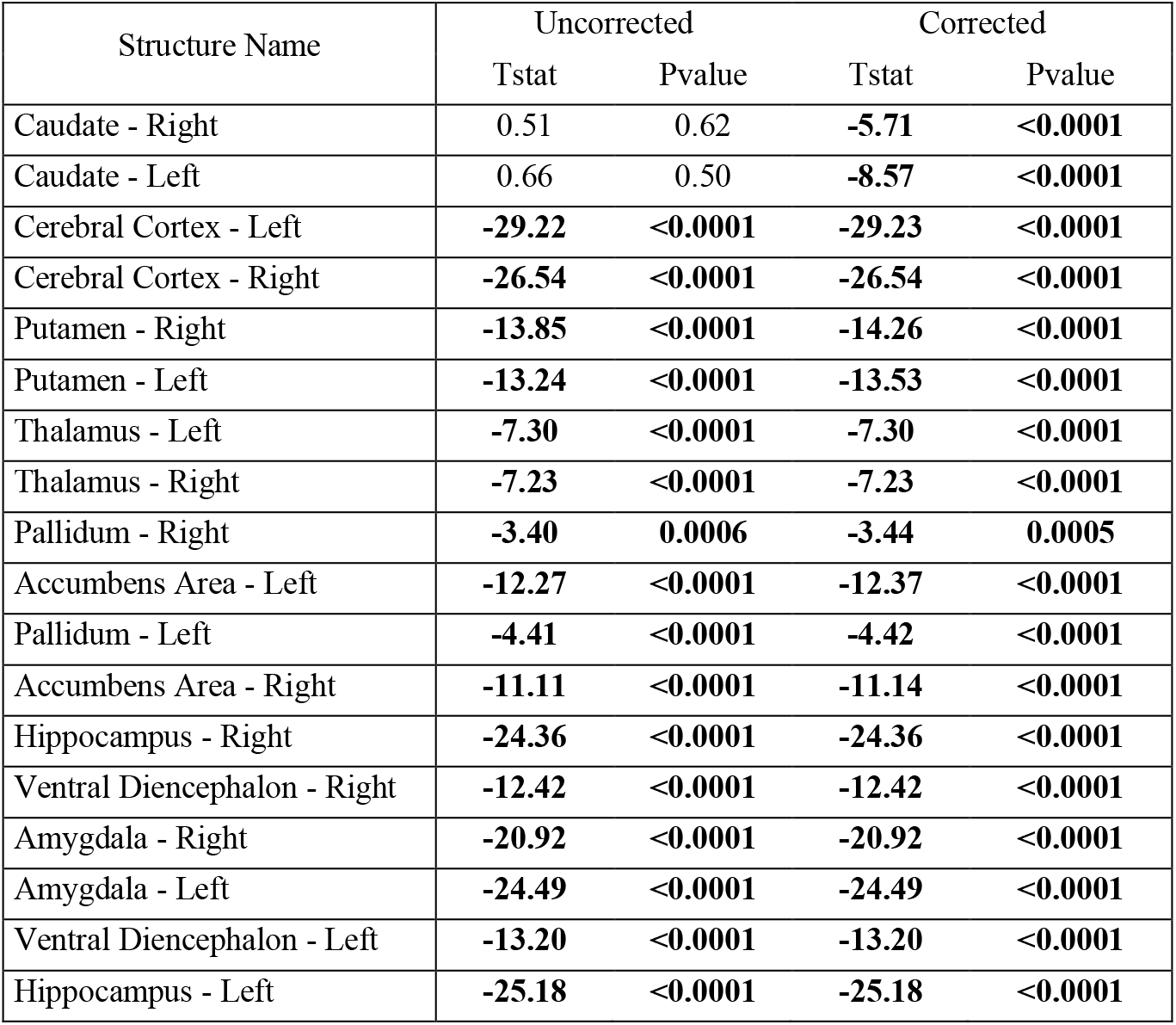
Associations between uncorrected and corrected GM volumes and ADAS13. Significant results after FDR correction are indicated in bold font.

| Structure Name | Uncorrected |  | Corrected |  |
| --- | --- | --- | --- | --- |
|  | Tstat | Pvalue | Tstat | Pvalue |
| Caudate - Right | 0.51 | 0.62 | **-5.71** | **<0.0001** |
| Caudate - Left | 0.66 | 0.50 | **-8.57** | **<0.0001** |
| Cerebral Cortex - Left | **-29.22** | **<0.0001** | **-29.23** | **<0.0001** |
| Cerebral Cortex - Right | **-26.54** | **<0.0001** | **-26.54** | **<0.0001** |
| Putamen - Right | **-13.85** | **<0.0001** | **-14.26** | **<0.0001** |
| Putamen - Left | **-13.24** | **<0.0001** | **-13.53** | **<0.0001** |
| Thalamus - Left | **-7.30** | **<0.0001** | **-7.30** | **<0.0001** |
| Thalamus - Right | **-7.23** | **<0.0001** | **-7.23** | **<0.0001** |
| Pallidum - Right | **-3.40** | **0.0006** | **-3.44** | **0.0005** |
| Accumbens Area - Left | **-12.27** | **<0.0001** | **-12.37** | **<0.0001** |
| Pallidum - Left | **-4.41** | **<0.0001** | **-4.42** | **<0.0001** |
| Accumbens Area - Right | **-11.11** | **<0.0001** | **-11.14** | **<0.0001** |
| Hippocampus - Right | **-24.36** | **<0.0001** | **-24.36** | **<0.0001** |
| Ventral Diencephalon - Right | **-12.42** | **<0.0001** | **-12.42** | **<0.0001** |
| Amygdala - Right | **-20.92** | **<0.0001** | **-20.92** | **<0.0001** |
| Amygdala - Left | **-24.49** | **<0.0001** | **-24.49** | **<0.0001** |
| Ventral Diencephalon - Left | **-13.20** | **<0.0001** | **-13.20** | **<0.0001** |
| Hippocampus - Left | **-25.18** | **<0.0001** | **-25.18** | **<0.0001** |

**Figure 5.**
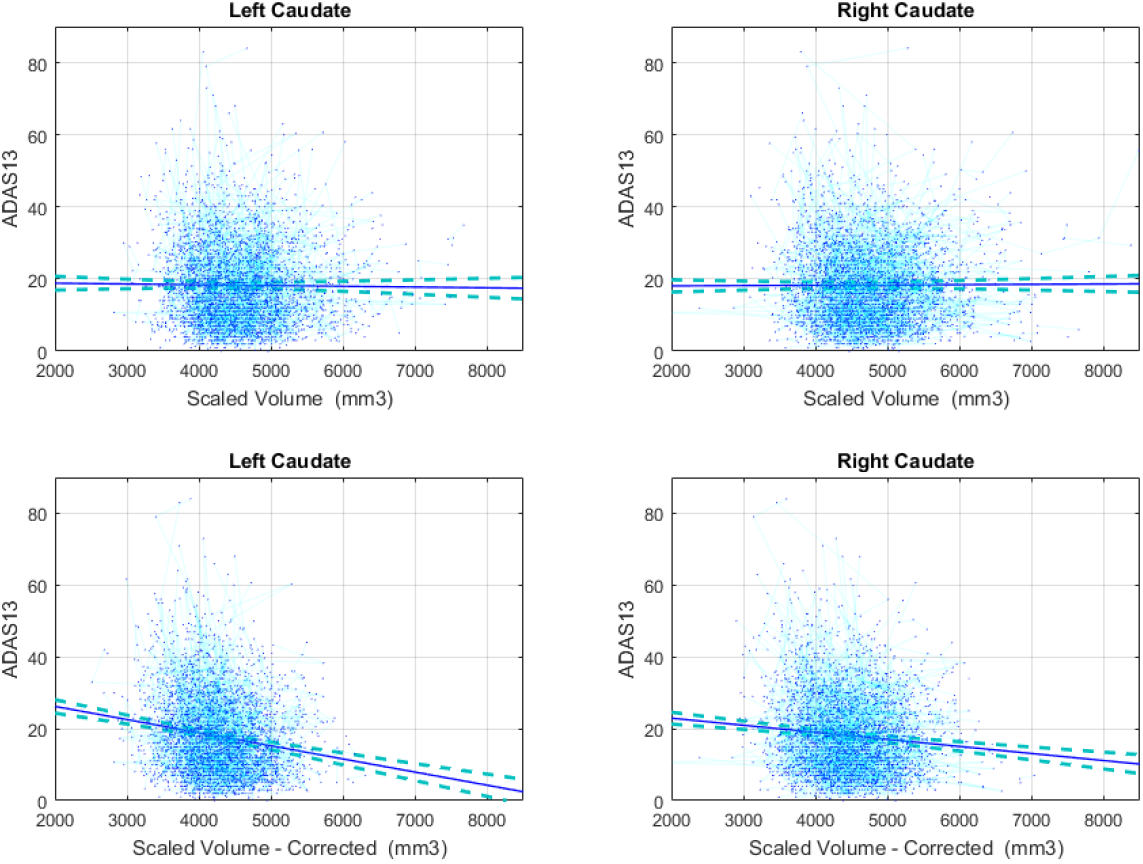
The association between uncorrected and corrected Caudate volumes ADAS13 scores.

## DISCUSSION

In this study, we investigated the impact of WMHs on GM segmentations, specifically for the *Freesurfer* segmentation tool, in order to determine whether WMHs might lead to systematic segmentation errors in certain GM regions, and whether such errors might impact associations between GM volumes and clinical outcomes. We found such errors in a number of regions, and in the caudate in particular it propagated to the association with clinical variables.

We found that overlapping voxel volumes between WMH and GM segmentations were significantly associated with overal WMH burden (Table 2), indicating higher error rates for subjects with high WMH loads. This affected both cortical and subcortical structures, and in particular the caudates bilaterally. Uncorrected, caudate volumes showed a significant *increase* with age, which is highly unlikely to be a real effect given that all regions, except the caudate, have been shown to decline in late-life cognitively healthy individuals (Potvin et al., 2016b, 2016a). This was further improbable given that the population of this study consists of not only aging individuals but also patients with MCI and AD, which are known to have increasing levels of atrophy across cortical areas. In contrast, the corrected caudate volumes significantly decreased with age, as could be expected. Given that WMHs are highly prevalent in the periventricular regions (i.e. white matter areas surrounding the caudate), the significant increase estimate is likely due to the fact that WMH burden increases with age and AD and MCI patients tend to have higher WMH loads and faster WMH progression.

Along the same line, corrected caudate volumes showed significant differences between AD and cognitively healthy cohorts, as well as significant associations with cognitive performance in the expected directions, whereas the uncorrected volumes were not significantly different and in the opposite direction. This again highlights the fact that if uncorrected for overlapping WMHs, estimates of caudate volumes can lead to incorrect or unreliable findings, particularly in populations with high prevalence of WMHs.

While other GM regions also had overlaps with WMHs that were significantly associated with the overal WMH burden (e.g. cerebral cortex, putamen), the amount and percentage of these overlaps were not nearly as high as caudate (17, 22, 80, and 74 mm^3^ versus 217 and 223 mm^3^ or 0.4%, 0.5%, 0.04%, and 0.04% versus 6.1% and 6.5%), and therefore, they did not affect their overall estimates and associations with age, diagnosis, and ADAS13. However, this might not be the case in other populations, where WMHs are more prevalent, or have different distributions.

*FreeSurfer* is one of the most widely used publicly available brain segmentation tools. Large databases such as the UK Biobank provide GM structure volumes derived from *FreeSurfer* to researchers. In line with our findings, other researchers have reported a positive association between WMH load and *FreeSurfer* caudate volumes in the UK biobank participants (Morys et al., 2020) and in another large sample of cognitively healthy individuals (Potvin et al., 2017a, 2017b). Studies investigating a larger age range report a U-shape curve for caudate volumes, decreasing from early adulthood to the 60s, and then increasing afterwards (Fjell et al., 2013, 2009; Goodro et al., 2012; Pfefferbaum et al., 2013; Potvin et al., 2016b; Walhovd et al., 2011). Given that WMHs generally occur in this same age range (i.e. after 60s), these results are also likely due to the segmentation errors caused by presence of WMHs in older participants.

Specifically for this algorithm, our study emphasizes the need for correcting *FreeSurfer* GM volume estimates for WMHs, particularly for the caudate. However, it is likely that other algorithms exhibit the same behavior. Evidence can be found in the literature, for example in the works by Goodro et al. and Pfefferbaum et al. using FSL volumes (Goodro et al., 2012; Pfefferbaum et al., 2013). Developers and users alike should therefore be aware of the possibility of systematic bias from not taking into account WMHs in GM segmentation from T1w images.

In conclusion, the presence of WMHs can lead to systematic errors in GM segmentations in certain regions, particularly in the caudate, which, if not corrected, can impact findings in populations with high WMH prevalence.

## Acknowledgements

MD is supported by a scholarship from the Canadian Consortium on Neurodegeneration in Aging in which SD and RC are co-investigators. The Consortium is supported by a grant from the Canadian Institutes of Health Research with funding from several partners including the Alzheimer Society of Canada, Sanofi, and Women’s Brain Health Initiative.

Data collection and sharing for this project was funded by the Alzheimer’s Disease Neuroimaging Initiative (ADNI) (National Institutes of Health Grant U01 AG024904) and DOD ADNI (Department of Defense award number W81XWH-12-2-0012). ADNI is funded by the National Institute on Aging, the National Institute of Biomedical Imaging and Bioengineering, and through generous contributions from the following: AbbVie, Alzheimer’s Association; Alzheimer’s Drug Discovery Foundation; Araclon Biotech; BioClinica, Inc.; Biogen; Bristol-Myers Squibb Company; CereSpir, Inc.; Cogstate; Eisai Inc.; Elan Pharmaceuticals, Inc.; Eli Lilly and Company; EuroImmun; F. Hoffmann-La Roche Ltd and its affiliated company Genentech, Inc.; Fujirebio; GE Healthcare; IXICO Ltd.; Janssen Alzheimer Immunotherapy Research & Development, LLC.; Johnson & Johnson Pharmaceutical Research & Development LLC.; Lumosity; Lundbeck; Merck & Co., Inc.; Meso Scale Diagnostics, LLC.; NeuroRx Research; Neurotrack Technologies; Novartis Pharmaceuticals Corporation; Pfizer Inc.; Piramal Imaging; Servier; Takeda Pharmaceutical Company; and Transition Therapeutics. The Canadian Institutes of Health Research is providing funds to support ADNI clinical sites in Canada. Private sector contributions are facilitated by the Foundation for the National Institutes of Health (www.fnih.org). The grantee organization is the Northern California Institute for Research and Education, and the study is coordinated by the Alzheimer’s Therapeutic Research Institute at the University of Southern California. ADNI data are disseminated by the Laboratory for Neuro Imaging at the University of Southern California.

## References

Abraham, H.M.A., Wolfson, L., Moscufo, N., Guttmann, C.R., Kaplan, R.F., White, W.B., 2016. Cardiovascular risk factors and small vessel disease of the brain: blood pressure, white matter lesions, and functional decline in older persons. J. Cereb. Blood Flow Metab. 36, 132–142.

Appelman, A.P., Exalto, L.G., Van Der Graaf, Y., Biessels, G.J., Mali, W.P., Geerlings, M.I., 2009. White matter lesions and brain atrophy: more than shared risk factors? A systematic review. Cerebrovasc. Dis. 28, 227–242.

Barber, R., Scheltens, P., Gholkar, A., Ballard, C., McKeith, I., Ince, P., Perry, R., O’Brien, J., 1999. White matter lesions on magnetic resonance imaging in dementia with Lewy bodies, Alzheimer’s disease, vascular dementia, and normal aging. J. Neurol. Neurosurg. Psychiatry 67, 66–72. https://doi.org/10.1136/jnnp.67.1.66

Benjamini, Y., Hochberg, Y., 1995. Controlling the false discovery rate: a practical and powerful approach to multiple testing. J. R. Stat. Soc. Ser. B Methodol. 57, 289–300.

Caroppo, P., Ber, I.L., Camuzat, A., Clot, F., Naccache, L., Lamari, F., Septenville, A.D., Bertrand, A., Belliard, S., Hannequin, D., Colliot, O., Brice, A., 2014. Extensive White Matter Involvement in Patients With Frontotemporal Lobar Degeneration: Think Progranulin. JAMA Neurol. 71, 1562–1566. https://doi.org/10.1001/jamaneurol.2014.1316

Dadar, M., Fonov, V.S., Collins, D.L., Initiative, A.D.N., 2018a. A comparison of publicly available linear MRI stereotaxic registration techniques. NeuroImage 174, 191–200.

Dadar, M., Maranzano, J., Ducharme, S., Carmichael, O.T., Decarli, C., Collins, D.L., 2017a. Validation of T1w-based segmentations of white matter hyperintensity volumes in large-scale datasets of aging. Hum. Brain Mapp.

Dadar, M., Maranzano, J., Ducharme, S., Carmichael, O.T., Decarli, C., Collins, D.L., Initiative, A.D.N., 2018b. Validation of T 1w-based segmentations of white matter hyperintensity volumes in large-scale datasets of aging. Hum. Brain Mapp. 39, 1093–1107.

Dadar, M., Maranzano, J., Ducharme, S., Collins, D.L., 2019. White matter in different regions evolves differently during progression to dementia. Neurobiol. Aging 76, 71–79. https://doi.org/10.1016/j.neurobiolaging.2018.12.004

Dadar, M., Maranzano, J., Misquitta, K., Anor, C.J., Fonov, V.S., Tartaglia, M.C., Carmichael, O.T., Decarli, C., Collins, D.L., Alzheimer’s Disease Neuroimaging Initiative, 2017b. Performance comparison of 10 different classification techniques in segmenting white matter hyperintensities in aging. NeuroImage 157, 233–249. https://doi.org/10.1016/j.neuroimage.2017.06.009

Dadar, M., Pascoal, T., Manitsirikul, S., Misquitta, K., Tartaglia, C., Brietner, J., Rosa-Neto, P., Carmichael, O., DeCarli, C., Collins, D.L., 2017c. Validation of a Regression Technique for Segmentation of White Matter Hyperintensities in Alzheimer’s Disease. IEEE Trans. Med. Imaging.

Debette, S., Markus, H.S., 2010. The clinical importance of white matter hyperintensities on brain magnetic resonance imaging: systematic review and meta-analysis. Bmj 341, c3666.

Fischl, B., 2012. FreeSurfer. Neuroimage 62, 774–781.

Fjell, A.M., Westlye, L.T., Amlien, I., Espeseth, T., Reinvang, I., Raz, N., Agartz, I., Salat, D.H., Greve, D.N., Fischl, B., Dale, A.M., Walhovd, K.B., 2009. Minute Effects of Sex on the Aging Brain: A Multisample Magnetic Resonance Imaging Study of Healthy Aging and Alzheimer’s Disease. J. Neurosci. 29, 8774–8783. https://doi.org/10.1523/JNEUROSCI.0115-09.2009

Fjell, A.M., Westlye, L.T., Grydeland, H., Amlien, I., Espeseth, T., Reinvang, I., Raz, N., Holland, D., Dale, A.M., Walhovd, K.B., 2013. Critical ages in the life course of the adult brain: nonlinear subcortical aging. Neurobiol. Aging 34, 2239–2247. https://doi.org/10.1016/j.neurobiolaging.2013.04.006

Goodro, M., Sameti, M., Patenaude, B., Fein, G., 2012. Age effect on subcortical structures in healthy adults. Psychiatry Res. Neuroimaging 203, 38–45.

Gouw, A.A., van der Flier, W.M., Fazekas, F., van Straaten, E.C., Pantoni, L., Poggesi, A., Inzitari, D., Erkinjuntti, T., Wahlund, L.O., Waldemar, G., 2008. Progression of white matter hyperintensities and incidence of new lacunes over a 3-year period: the Leukoaraiosis and Disability study. Stroke 39, 1414–1420.

Kandiah, N., Mak, E., Ng, A., Huang, S., Au, W.L., Sitoh, Y.Y., Tan, L.C.S., 2013. Cerebral white matter hyperintensity in Parkinson’s disease: A major risk factor for mild cognitive impairment. Parkinsonism Relat. Disord. 19, 680–683. https://doi.org/10.1016/j.parkreldis.2013.03.008

Manjón, J.V., Coupé, P., Martí-Bonmatí, L., Collins, D.L., Robles, M., 2010. Adaptive non-local means denoising of MR images with spatially varying noise levels. J. Magn. Reson. Imaging 31, 192–203.

Mateos-Pérez, J.M., Dadar, M., Lacalle-Aurioles, M., Iturria-Medina, Y., Zeighami, Y., Evans, A.C., 2018. Structural neuroimaging as clinical predictor: A review of machine learning applications. NeuroImage Clin. https://doi.org/10.1016/j.nicl.2018.08.019

Mohs, R.C., Cohen, L., 1987. Alzheimer’s Disease Assessment Scale (ADAS). Psychopharmacol. Bull. 24, 627–628.

Morys, F., Dadar, M., Dagher, A., 2020. Obesity impairs cognitive function via metabolic syndrome and cerebrovascular disease: an SEM analysis in 15,000 adults from the UK Biobank. bioRxiv.

Petersen, R.C., Aisen, P.S., Beckett, L.A., Donohue, M.C., Gamst, A.C., Harvey, D.J., Jack, C.R., Jagust, W.J., Shaw, L.M., Toga, A.W., others, 2010. Alzheimer’s disease Neuroimaging Initiative (ADNI) clinical characterization. Neurology 74, 201–209.

Pfefferbaum, A., Rohlfing, T., Rosenbloom, M.J., Chu, W., Colrain, I.M., Sullivan, E.V., 2013. Variation in longitudinal trajectories of regional brain volumes of healthy men and women (ages 10 to 85 years) measured with atlas-based parcellation of MRI. Neuroimage 65, 176–193.

Potvin, O., Dieumegarde, L., Duchesne, S., Initiative, A.D.N., 2017a. Normative morphometric data for cerebral cortical areas over the lifetime of the adult human brain. Neuroimage 156, 315–339.

Potvin, O., Dieumegarde, L., Duchesne, S., Initiative, A.D.N., 2017b. Normative morphometric data for cerebral cortical areas over the lifetime of the adult human brain. Neuroimage 156, 315–339.

Potvin, O., Mouiha, A., Dieumegarde, L., Duchesne, S., Initiative, A. s D.N., 2016a. FreeSurfer subcortical normative data. Data Brief 9, 732.

Potvin, O., Mouiha, A., Dieumegarde, L., Duchesne, S., Initiative, A.D.N., 2016b. Normative data for subcortical regional volumes over the lifetime of the adult human brain. Neuroimage 137, 9–20.

Raman, M.R., Kantarci, K., Murray, M.E., Jack, C.R., Vemuri, P., 2016. Imaging markers of cerebrovascular pathologies: Pathophysiology, clinical presentation, and risk factors. Alzheimers Dement. Diagn. Assess. Dis. Monit. 5, 5–14. https://doi.org/10.1016/j.dadm.2016.12.006

Sudre, C.H., Bocchetta, M., Cash, D., Thomas, D.L., Woollacott, I., Dick, K.M., van Swieten, J., Borroni, B., Galimberti, D., Masellis, M., Tartaglia, M.C., Rowe, J.B., Graff, C., Tagliavini, F., Frisoni, G., Laforce, R., Finger, E., de Mendonça, A., Sorbi, S., Ourselin, S., Cardoso, M.J., Rohrer, J.D., Andersson, C., Archetti, S., Arighi, A., Benussi, L., Binetti, G., Black, S., Cosseddu, M., Fallström, M., Ferreira, C., Fenoglio, C., Fox, N.C., Freedman, M., Fumagalli, G., Gazzina, S., Ghidoni, R., Grisoli, M., Jelic, V., Jiskoot, L., Keren, R., Lombardi, G., Maruta, C., Mead, S., Meeter, L., van Minkelen, R., Nacmias, B., Öijerstedt, L., Padovani, A., Panman, J., Pievani, M., Polito, C., Premi, E., Prioni, S., Rademakers, R., Redaelli, V., Rogaeva, E., Rossi, G., Rossor, M.N., Scarpini, E., Tang-Wai, D., Thonberg, H., Tiraboschi, P., Verdelho, A., Warren, J.D., 2017. White matter hyperintensities are seen only in GRN mutation carriers in the GENFI cohort. NeuroImage Clin. 15, 171–180. https://doi.org/10.1016/j.nicl.2017.04.015

Walhovd, K.B., Westlye, L.T., Amlien, I., Espeseth, T., Reinvang, I., Raz, N., Agartz, I., Salat, D.H., Greve, D.N., Fischl, B., Dale, A.M., Fjell, A.M., 2011. Consistent neuroanatomical age-related volume differences across multiple samples. Neurobiol. Aging 32, 916–932. https://doi.org/10.1016/j.neurobiolaging.2009.05.013

